# Serotyping *Salmonella* Enteritidis and Typhimurium Using Whole Cell Matrix Assisted Laser Desorption Ionization – Time of Flight Mass Spectrometry (MALDI-TOF MS) through Multivariate Analysis and Artificial Intelligence

**DOI:** 10.1101/2022.07.01.498527

**Authors:** Anli Gao, Jennifer Fischer-Jenssen, Durda Slavic, Kimani Rutherford, Sarah Lippert, Emily Wilson, Shu Chen, Carlos G. Leon-Velarde, Perry Martos

## Abstract

*Salmonella* is one of the most frequent food-borne zoonoses, while *Salmonella* Typhimurium and Enteritidis are the major serovars of concern in public health. 113 *Salmonella* strains including 38 *S*. Enteritidis (SE), 38 *S*. Typhimurium (ST) and 37 strains of 32 other *Salmonella* serovars (SG) were tested in quadruplicate by whole-cell MALDI-TOF MS. Ions were studied and aligned from the raw data of mzXML files using Mass-Up (http://www.sing-group.org/mass-up), resulting in 1,741 aligned peaks. Datasets of ions (presence/absence) selected using a home-developed criteria on their specificity and detectability were subjected to multivariate analyses and artificial intelligence tools. Principle Component Analysis based on 88 selected ions separated SE, ST and SG without overlap on the first three principle components. The network and forest based deep learning tools were more sophisticated than the decision tree-based models. Neural Network carried out consistently well in training model, but no advantage was gained over the other models in validation results. HP (high performance) Neural, Support Vector Machine, HP Forest and Gradient Boasting were able to identify SE, ST and SG up to 100% correctly in both training and validation when 88 selected ions were used in analysis. Among them, HP Neural seemed to perform slightly better and relatively stable. Selection of serovar specific ions helps develop serotyping by increasing signal to noise. MALDI-TOF MS used with appropriate data processing strategies and classification tools could be applied to quickly alert when *Salmonella* serotypes of concern are suspected among routinely processed samples.

## Introduction

*Salmonella* is the etiologic agent of Salmonellosis in both humans and poultry capable of causing severe enteric illness with varying outcomes, particularly in the elderly, infants and immunocompromised individuals. Most infections develop from the ingestion of foods of animal origin contaminated with *Salmonella* species, with *Salmonella* Enteritidis (SE) and Typhimurium (ST) considered major serovars of concern in public health (1, 2). Rapid typing has been an important tool in identifying the pathogenic strains involved in an outbreak, in order to trace the potential source of infection and assess the risk (3). The traditional White-Kauffmann-Le Minor serotyping method is based on a combination of immunological reactions with the somatic O, flagellar H, and capsular Vi antigens (4). This method is time-consuming, and requires >250 different antisera to differentiate over 2600 serovars (5). Interpreting the aggregate results requires considerable expertise. While PCR, whole-genome sequencing and other molecular methods are useful for typing (6, 7, 8), the identification of microorganisms using Matrix-Assisted Laser Desorption Ionization Time-Of-Flight Mass Spectrometry (MALDI-TOF MS) has been accepted in clinical diagnosis at species level identification for its high-volume capacity, low running cost and rapid, reliable results that are simple to interpret (9). Significant effort has been made in subspecies typing using MALDI-TOF MS for a number of bacterial species including *Salmonella* with mixed claims, such as suitable, unsuitable, or unable to conclude (5, 10–18).

The aim of this study was to investigate the utility of multivariate analysis and artificial intelligence (AI) applied to the raw data of *Salmonella* MALDI-TOF MS spectra in order to identify SE and ST strains. First, MALDI-TOF MS spectra were analyzed with optimized peak picking and alignment protocol. Second, datasets which included ion peaks with relatively high discriminative power were created and subjected to classification using both a linear model and AI tools. Third, the outcome from different models were compared along with factors that affected the typing results.

## Materials and Methods

### Bacterial strains

A total of 113 *Salmonella* strains were assigned into three serovar groups (Table 1). Group A consisted of 38 SE strains; Group B 38 ST strains and Group C 37 *Salmonella* strains belong to 32 other serovars, hereafter referred to as *Salmonella* generic (SG) strains. Isolates were obtained from the American Type Culture Collection (ATCC, Manassas, Virginia, USA), National Microbiology Laboratory at Guelph (Guelph, ON, Canada) or isolated from routine analysis of poultry environments or food samples spanning 20 years. All strains except ATCC strains were serotyped by the National Microbiology Laboratory at Guelph (Guelph, ON, Canada). The O or somatic antigens of the *Salmonella* isolates were determined by slide agglutination (19). The H or flagellar antigens were identified using a microtechnique that employs micro-titre plates (20). The serovars were named using the antigenic formulae (21). All strains were retrieved from −80° C storage stock. Single colonies were isolated on Mueller Hinton agar supplemented with 5% sheep blood (MHBA) and incubated at 35 °C for 18-24 h. Five batches of samples were analyzed using MALDI-TOF MS. Mixed strains from different groups in a batch were studied, which minimized the effect of batch performance on the classification of serotypes (Table 1).

**Table 1.**
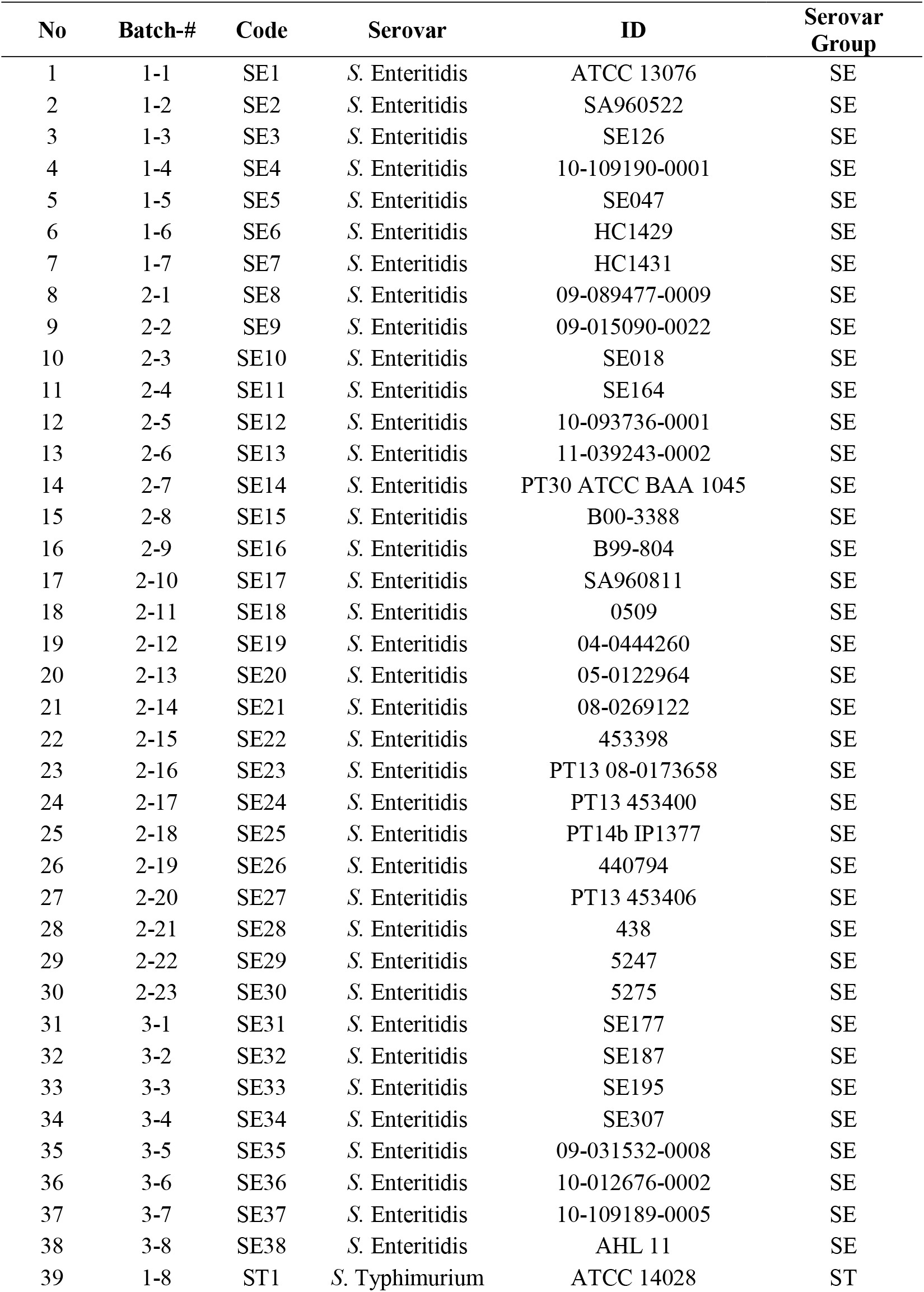
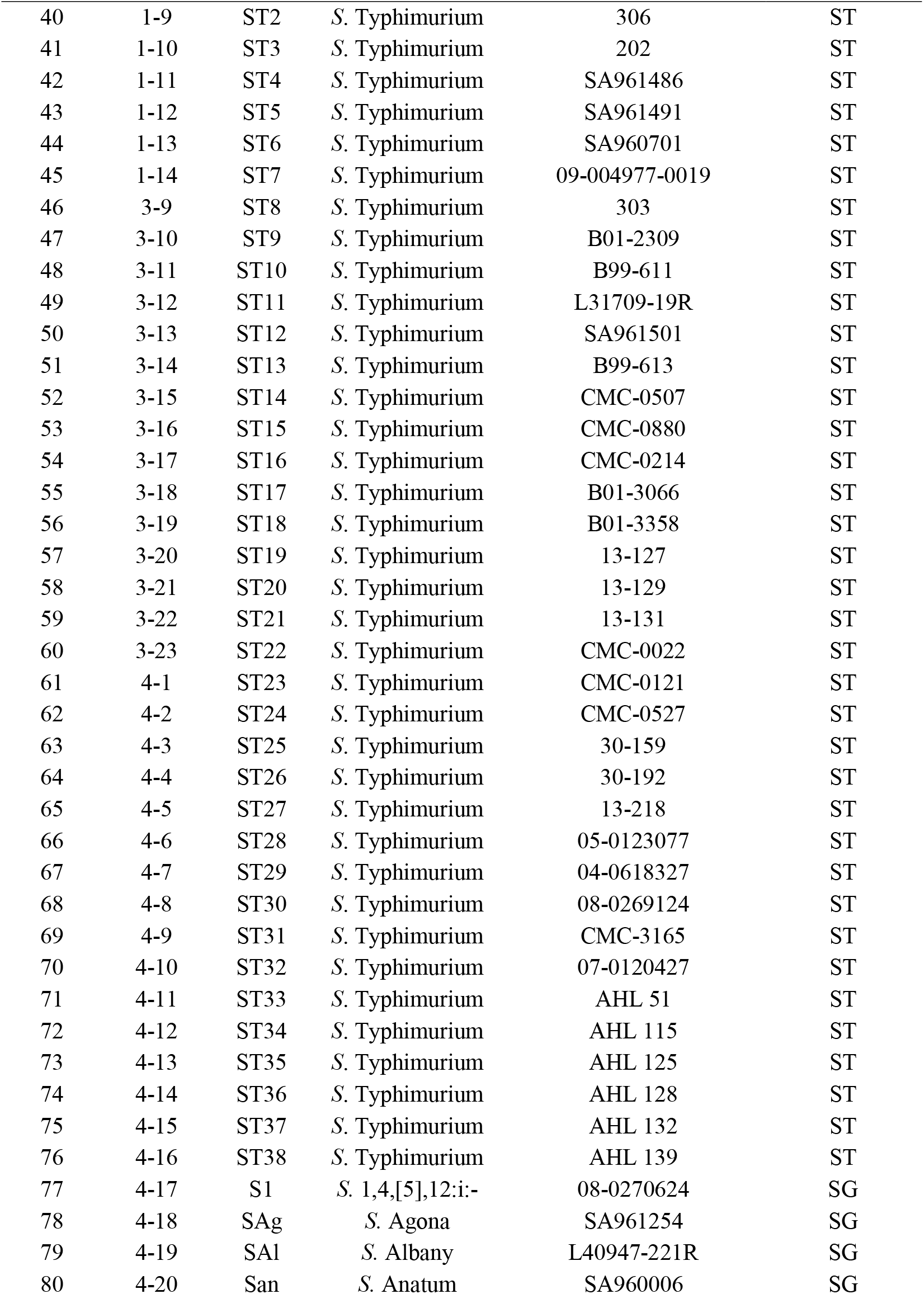
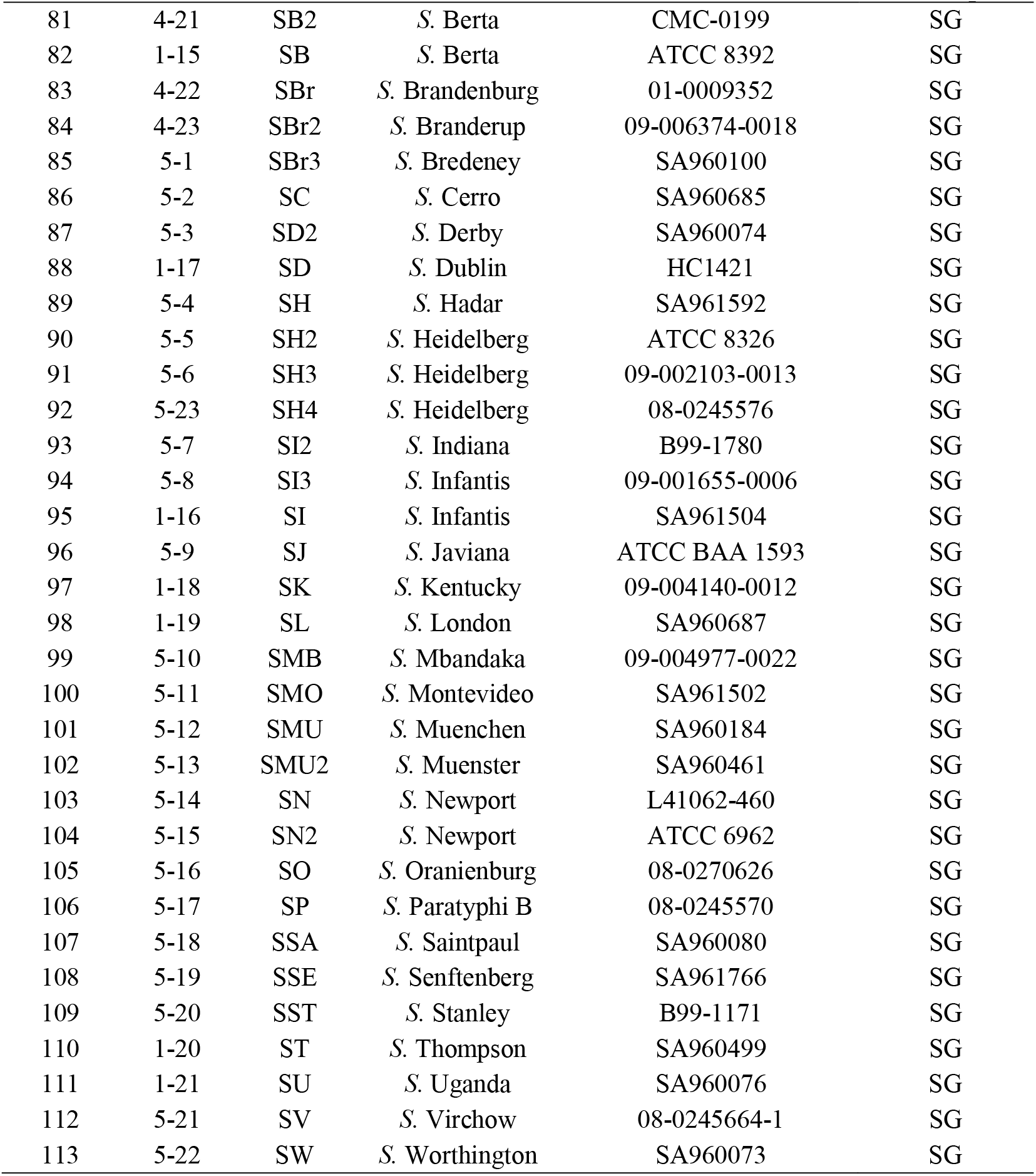
*Salmonella* strains analyzed using MALDI TOF MS for differentiation of *S*. Enteritidis and *S*. Typhimurium.

### Whole-cell MALDI-TOF MS

A small portion of bacterial colony was collected from the MHBA plate using a sterile 1 µL inoculating loop and smeared onto the stainless steel target plate (n = 4). Each bacterial smear was coated with 1 µL of α-cyano-hydroxycinnamic acid (HCCA) matrix and left to air dry. Mass spectra were acquired with a MicroFlex Biotyper LT System (Bruker Daltonik GmbH, Bremen, Germany) equipped with an all-solid-state Smartbeam Nd:YAG laser and operated at 100 Hz in the positive linear mode. Each spectrum was obtained by averaging automatic acquisition of 240 laser shots. A commercially available *E. coli* control was used with each batch of samples for internal calibration.

### Data analysis

Raw spectra in mzXML format with 2,000 – 21,000 m/z were acquired using the Export function in FlexAnalysis, organized in three subdirectories corresponding to the three serovar groups, SE, ST and SG, and analyzed using MASS-UP (22, http://www.sing-group.org/mass-up/) for peak detection and matching for alignment. The spectra in the three directories were loaded in labeled type, resulting in three folders, SE, ST, and SG in “Labeled Raw Data”, and processed with an optimized protocol in detection of peaks with the following settings, Intensity transformation: Log; Smoothing: Savitzky Golay; Baseline correction: SNIP (Statistics-sensitive Non-linear Iterative Peak-clipping algorithm); Standardization: Probabilistic Quotient Normalization; Peak detection: MSW (SNR: 6; PakScaleRange: 2; amp.th: 0.0001) for peak identification. Then, peak matching was carried out using MALDIquant (Tolerance type: PPM; Tolerance: 300; and Reference type: AVG) for both intra-sample and inter-sample matching at Tolerance = 0.002 with consensus spectrum at Percentage of presence of 50%. After the analysis of Biomarker discovery (intra class analysis) [Labeled], the intensity and the presence/absence of matched peaks of all tested samples were passed and integrated in Excel for further analysis. The number of detected peaks ranged from 360 – 450 per test in 452 individual tests. The final dataset contained 1741 aligned ions from 1995 – 20075 m/z, including 39 unary peaks repeatedly detected in all of the 113 strains (Table 3).

Principle Component Analysis (PCA) and Cluster Analysis were carried out with Unscrambler X 10.5 (CAMO Software AS, Oslo, Norway). Seven principle components were calculated with cross validation conducted randomly in 20 segments, 5-6 samples per segment. Correlation was used as the measurement of distance and Hierarchical Average-link for merging of clusters in Cluster Analysis.

AI tools, such as HP (high performance) Tree, Decision Tree, Neural Network, HP Neural, Support Vector Machine, HP Forest and Gradient Boosting were conducted using SAS Enterprise Miner 15.1 (SAS 9.4, SAS Institute Inc., Cary, NC). The dataset was partitioned using HP Data Partition function, such that 70% of the samples from each group (SE: 27; ST: 27; SG: 26) were used for training and the remaining 30% (11 from each) for model validation.

## Results

### Sorting of peaks for biomarker discovery

An in-house developed formula [1] was used to evaluate individual peaks, in order to select the top peaks with relatively high discriminative power (DP) between serotypes.

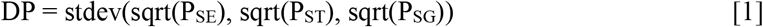

where P_SE_, percentage of the peak detected in SE; P_ST_, that in ST; P_SG_, that in SG. The stdev() and sqrt() are available functions in Excel. A large DP implies a higher discriminative power of a given peak for differentiating serovars. The sqrt() was adopted in the standard deviation function after it was discovered that stdev(sqrt(P_i_)) evaluated the peak presents solely in one serovar (an ideal biomarker) with higher DP value, while considering the detectability of the ion peaks as well, comparing with the function stdev(P_i_). Datasets containing different numbers of selected peaks with relatively high DP were generated, in order to evaluate their efficacy for subtyping using different analysis tools.

Table 2 presents the top 50 peaks with relatively high DP. A previously reported biomarker peak 3018 m/z by Yang *et al*. (16) was able to identify 31 out of 38 SE strains. Ions 6039 m/z, 5969 m/z and 3623 m/z identified in this study showed specificity for SE; however, these ion abundancies were relatively low, and information offered by these ions was mostly masked by 3018 m/z. A combination of ions 3018 m/z and 7640 m/z identified 36 SE strains out of 38 with 2 misclassifications, 1 as SG and 1 ST. In contrast, the ion 7100 m/z identified 31 out of 38 ST strains. When ion 20075 m/z, which identified 12 ST and 5 SG, was combined with 7100 m/z, 35 out of 38 ST strains were subsequently correctly identified with 3 misidentifications as SG. In general, the use of more potential biomarkers increases the sensitivity but decreases the specificity.

**Table 2.** Top 50 ion peaks, m/z, sorted based on their discriminative power (DP), a criterion developed in-house.

| No | Ion<br>(m/z) | SE<br>(N=38) | ST<br>(N=38) | SG<br>(N=37) | DP | No | Ion<br>(m/z) | SE<br>(N=38) | ST<br>(N=38) | SG<br>(N=37) | DP |
| --- | --- | --- | --- | --- | --- | --- | --- | --- | --- | --- | --- |
| 1 | 3018 | 31 | 0 | 0 | 0.52 | 26 | 6077 | 9 | 0 | 0 | 0.28 |
| 2 | 6039 | 30 | 0 | 0 | 0.51 | 27 | 10634 | 9 | 0 | 0 | 0.28 |
| 3 | 7100 | 0 | 31 | 3 | 0.46 | 28 | 6011 | 9 | 35 | 35 | 0.28 |
| 4 | 3550 | 0 | 30 | 2 | 0.46 | 29 | 2028 | 11 | 0 | 1 | 0.28 |
| 5 | 3254 | 28 | 0 | 12 | 0.44 | 30 | 9344 | 11 | 0 | 1 | 0.28 |
| 6 | 5969 | 21 | 0 | 0 | 0.43 | 31 | 3814 | 18 | 1 | 3 | 0.28 |
| 7 | 3623 | 17 | 0 | 0 | 0.39 | 32 | 4238 | 19 | 1 | 9 | 0.27 |
| 8 | 7640 | 25 | 1 | 1 | 0.37 | 33 | 6775 | 19 | 1 | 9 | 0.27 |
| 9 | 3004 | 3 | 37 | 26 | 0.37 | 34 | 12275 | 6 | 31 | 25 | 0.27 |
| 10 | 8820 | 2 | 31 | 26 | 0.37 | 35 | 5423 | 13 | 17 | 1 | 0.27 |
| 11 | 5781 | 20 | 2 | 0 | 0.37 | 36 | 15871 | 10 | 0 | 5 | 0.26 |
| 12 | 8800 | 20 | 0 | 4 | 0.36 | 37 | 8852 | 36 | 8 | 14 | 0.26 |
| 13 | 6498 | 0 | 14 | 0 | 0.35 | 38 | 3451 | 10 | 0 | 1 | 0.26 |
| 14 | 10883 | 4 | 0 | 17 | 0.34 | 39 | 12298 | 25 | 4 | 6 | 0.26 |
| 15 | 3700 | 31 | 2 | 14 | 0.34 | 40 | 7163 | 10 | 2 | 0 | 0.26 |
| 16 | 8747 | 9 | 0 | 15 | 0.33 | 41 | 4370 | 10 | 3 | 0 | 0.26 |
| 17 | 12533 | 12 | 0 | 0 | 0.32 | 42 | 3302 | 14 | 1 | 1 | 0.26 |
| 18 | 6657 | 14 | 0 | 6 | 0.31 | 43 | 9016 | 1 | 5 | 16 | 0.25 |
| 19 | 11414 | 13 | 4 | 0 | 0.29 | 44 | 14950 | 8 | 6 | 0 | 0.25 |
| 20 | 2998 | 25 | 2 | 12 | 0.29 | 45 | 2984 | 9 | 0 | 1 | 0.25 |
| 21 | 7071 | 21 | 1 | 8 | 0.29 | 46 | 12779 | 9 | 2 | 0 | 0.24 |
| 22 | 5312 | 2 | 4 | 22 | 0.29 | 47 | 13662 | 9 | 2 | 0 | 0.24 |
| 23 | 20075 | 0 | 12 | 5 | 0.29 | 48 | 12192 | 16 | 2 | 2 | 0.24 |
| 24 | 4953 | 27 | 3 | 9 | 0.28 | 49 | 5294 | 6 | 0 | 7 | 0.24 |
| 25 | 11486 | 12 | 4 | 0 | 0.28 | 50 | 7974 | 19 | 2 | 7 | 0.24 |

m/z, mass/charge; SE, *S. Enteritidis*; ST, *S. Typhimurium*; SG, other *Salmonella* serovars; DP, Discriminative power by the function described in “Sorting of peaks for biomarker discovery”.

**Table 3.** Unary ions (N = 39) identified in all of the 113 *Salmonella* strains (potentially useable for alignment of ion peaks from different researches).

|  |  |  |  |  |  |  |  |
| --- | --- | --- | --- | --- | --- | --- | --- |
| 2034 | 2573 | 3429 | 4462 | 5146 | 7118 | 8331 | 10069 |
| 2179 | 2616 | 3580 | 4499 | 5384 | 7264 | 8372 | 10288 |
| 2382 | 2689 | 3861 | 4622 | 5742 | 7325 | 8922 | 11216 |
| 2405 | 2825 | 4186 | 4765 | 6386 | 7722 | 8996 | 11239 |
| 2451 | 3127 | 4351 | 5036 | 6575 | 8122 | 9242 |  |

### Unsupervised analyses

PCA was carried out on both the quantitative dataset with intensity of peak signal and the qualitative dataset with presence (assigned to 1) / absence (assigned to 0) of every detected ion. Although the quantitative dataset had shown the power of differentiating among strains (unpublished data), it was not useful in the differentiation of serotypes. When quantitative data were analyzed the high abundance ions masked the low abundance ions. This is consistent with the findings of De Bruyne *et al*. (10) which showed that taking only the binary data into account seemed promising for the identification of *Leuconostoc* and *Fructobacillus* species.

Figure 1 presents the scatterplot of individual samples from the PCA of 88 peaks selected based on their DP. When the first 55 peaks were involved in analysis, touching but without overlap between SG and ST was observed. When the first 30 ions were used, overlap of a few samples between ST and SG was observed. The separation between serotypes was also reduced significantly when the first 222 peaks were studied (data not shown). Differences among serotypes were not observed when all 1,741 ions were included in the analyses. Although most of the SE, ST and SG samples preferred to merge within the same serovar group first in Cluster Analysis, early clustering of the samples from different groups was significant (data not shown), in agreement with literature (16).

**Figure 1.**
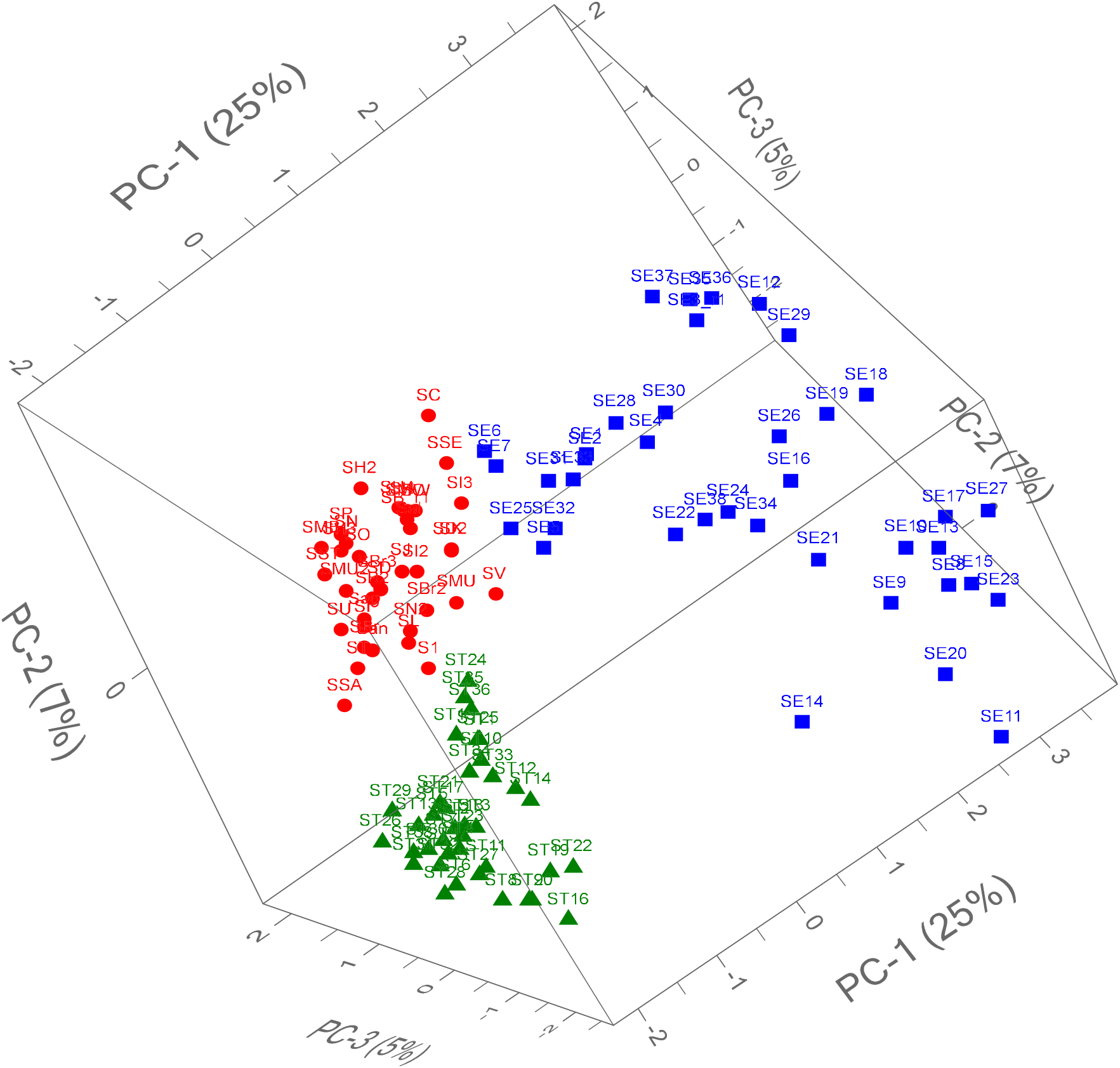
Principle component analysis (PCA) generated from 88 selected MALDI-TOF MS peaks with a validation of 20 segments. Green triangle, *S*. Typhimurium (N = 38); Blue square, *S*. Enteritidis (N = 38); Red dot: Mixed strains from 33 other *Salmonella* serovars (N = 37). Strain S1 (1,4,[5],12:i:-) is a monophasic variant of *S*. Typhimurium. Cf. Table 1 for more information on strains.

Serovar-unspecific ions introduce significant noise into the datasets, which dominated the signal from serovar specific peaks in PCA and subsequently reduced the discriminative power of subtyping. The profile representing a specific serovar could consist of a combination of peaks, in which some were strong and highly repeatable, while others were weak and not consistently detected (Table 2). Useful information could be lost when low abundance ions were not selected by the instrument and/or by the data processing protocols employed during analysis. For example, an ion selection procedure was not reported in the work by Kang et al. (12), and their conclusion that MALDI-TOF is of limited use to subtype *Salmonella* serovars could likely be reconsidered if a dataset containing serovar-specific peaks was used in the analysis.

Serovar-specific ions as biomarkers could be shared among serovars in *Salmonella* as shown in Table 2. The ion m/z may change due to substitution of amino acid and some serovars showed a notable degree of intra-serovar variability in MALDI-TOF spectra (5). As a result in MALDI-TOF MS, one mass peak could disappear while another appear. Therefore, a combination of biomarkers and sophisticated analytical tools are required for the identification of serotypes.

### Supervised Artificial intelligence tools

Machine learning methods handle multi-dimensional data easily and can offer solutions where standard analysis methods fail. Support Vector Machines and Random Forest have been employed and resulted in accuracies between 94% and 98% for the identification of *Leuconostoc* and *Fructobacillus* species, respectively (10). Figure 2 presents the misclassification rate (MR) of training and validation from the use of seven AI tools based on selected datasets with different partition strategies and seeds. Figures 2A and 2B represent the results of train and validation respectively with same partition strategy and seed. Figures 2C and 2D are from an analysis using different partition strategy and seed. Similar results from two additional analyses are not presented. Generally, the MR from train (Figures 2A and 2C) were lower than that from validation (Figures 2B and 2D) for a given model. The performance of HP Tree and Decision Tree was not affected apparently by the number of selected top biomarker peaks due to the nature of the models. The algorithm of these two models was programed to select the variables with top DP between supervised subgroups. Variables with relatively low DP below a criterion were not considered. Their MR, 10% - 20%, matches the proportion of the samples those miss the top biomarkers with high DP (Cf. Table 2).

**Figure 2.**
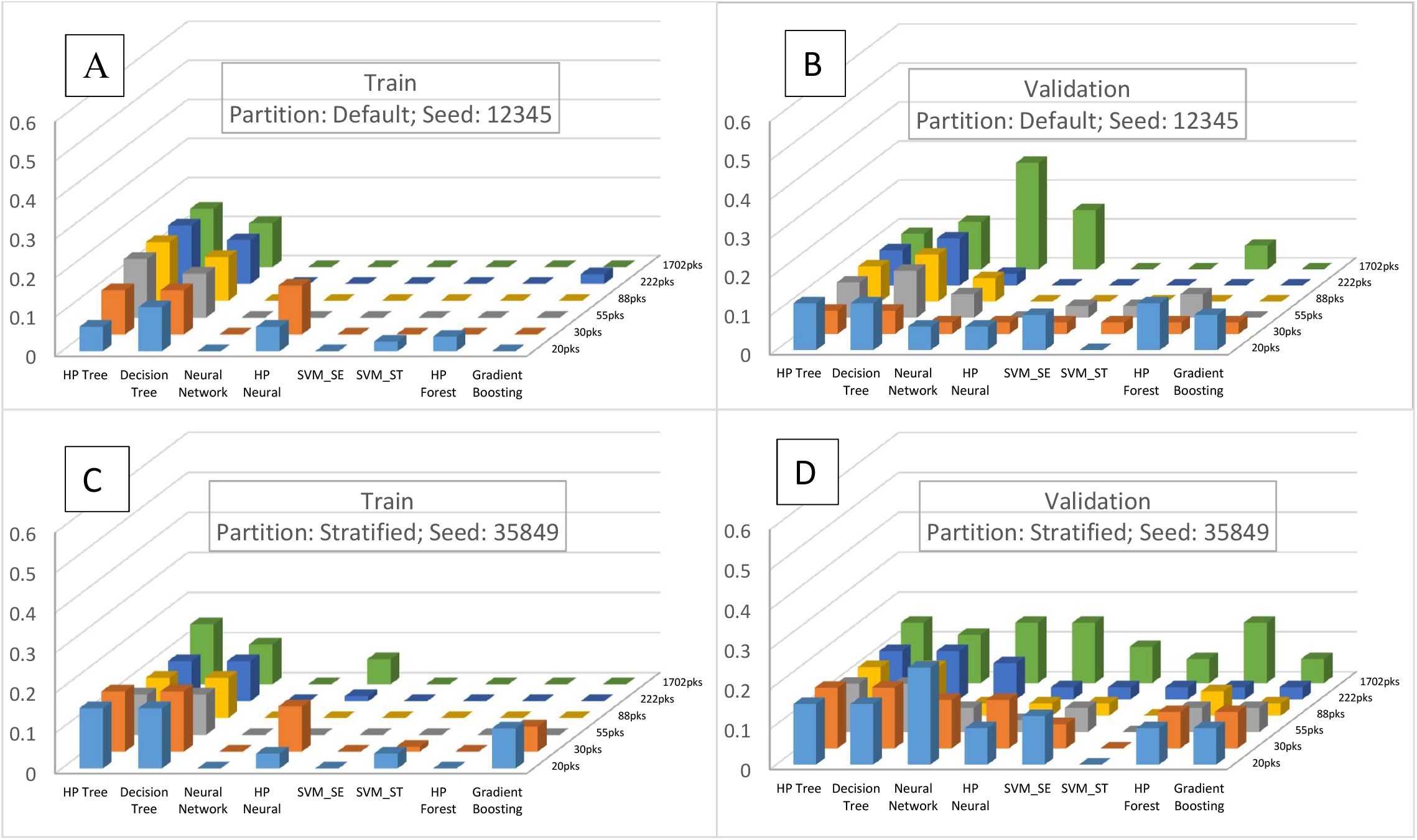
Misclassification rate (MR) of *Salmonella* serovars of train (70% of samples) and validation (30% of samples) in seven artificial intelligent tools (from left to right: HP Tree, Decision Tree, Neural Network, HP Neural, SVM (Support Vector Machine), HP Forest, and Gradient Boosting) based on datasets containing different number of peaks selected according to their discriminative power (Cf. Table 2). SVM_SE targets *S*. Enteritidis, while SVM_ST targets *S*. Typhimurium only. Note: A and B represent combination resulting in relatively low MR, whilst C and D represent that resulting in high MR.

Figure 3 illustrates the chart produced by Decision Tree based on 88 selected ions with partition method “default” and seed “12345”. In this model, 3018 m/z, the first top specific biomarker for SE (Table 2), was used to identify SE to Node 3. Then ion 7100 m/z was used in Node 2 to separate most of the ST including a few SG to Node 5. Furthermore, ion 7640 m/z was used in Node 4 to separate a few more SE out to Node 7. In the HP Tree, the only difference is that ion 20075 m/z was used in the place of ion 7640 m/z (p7640 in Figure 3) to separate some SG plus a few SE and one ST to Node 5 and a few ST and SG (without SE) to Node 6 (chart not shown). Ion 3018 m/z is SE-specific, but was not 100% repeatable in SE. Ion 7100 m/z, as well as 7640 and 20075 m/z, seemed discriminative, but was common with other serovar(s) (Table 2).

**Figure 3.**
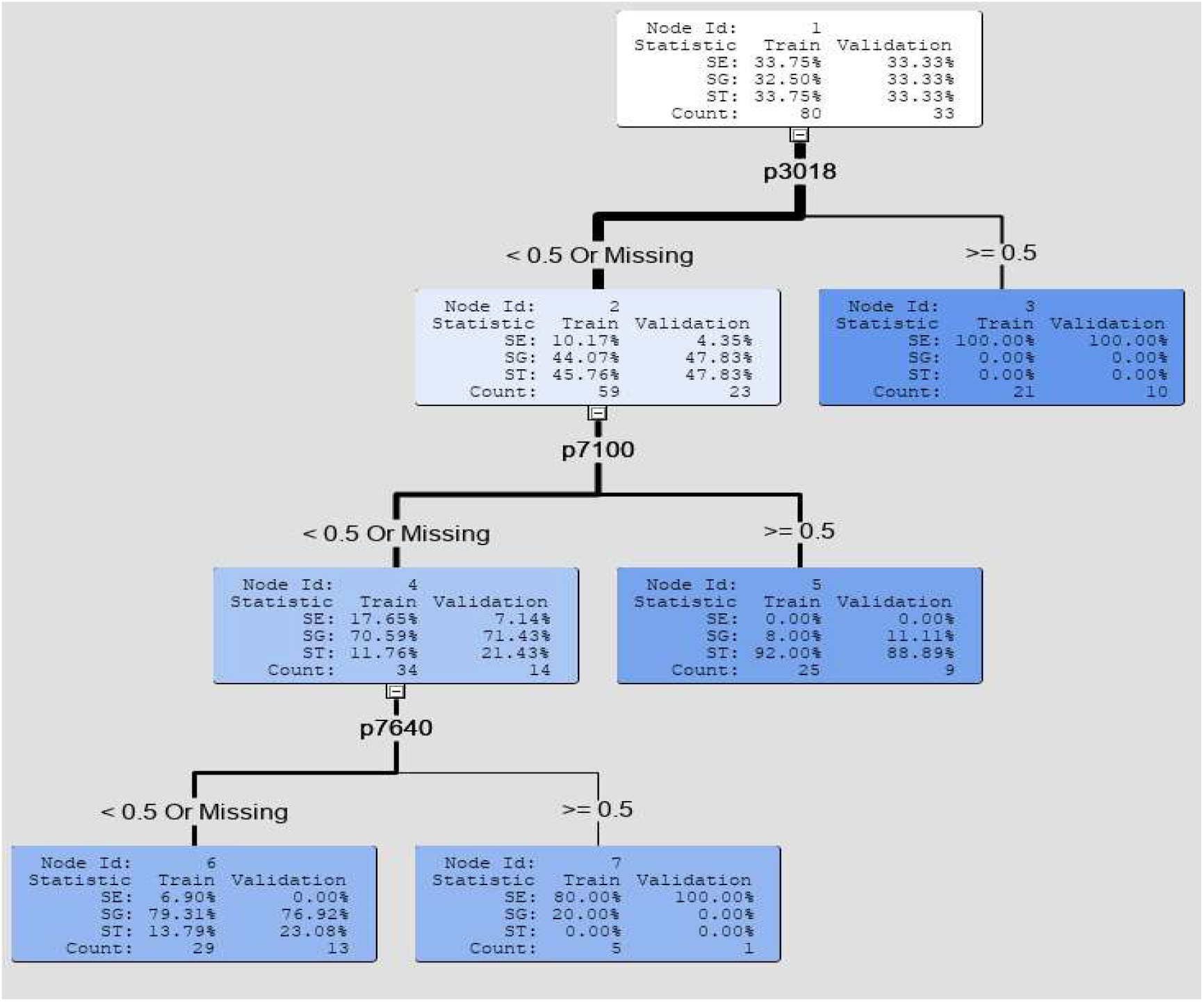
Diagram of Decision Tree based on 88 selected peaks in Figure 2A. Peak 20075 (p20075) was used in the place of peak 7640 (p7640) in the diagram of HP Tree. Cf. Figure 2 for comparison with other models. “<0.5 Or Missing” means “0 (absent)”, while “>=0.5” means “1 (present)” in this study.

Although Neural Network resulted in relatively low MR in training, the performance in validation was not favourable (Figure 2). This observation was consistent with reports where the Neural Network as a deep learning approach had an issue of overfitting of the training model due to the inclusion of noise (23). This emphasizes the necessity of model validation in model evaluation. The HP Neural is a derivative of Neural Network, but it provided better and more balanced results between training and validation compared to Neural Network on overcoming the overfitting issue. Forest is an ensemble of decision trees. Therefore, Forest brings the advent of decision making from a collection of decision trees (Cf. HP Forest against HP Tree and Decision Tree in Figure 2). Nevertheless, HP Neural seems to perform with slightly more stability than HP Forest and Gradient Boosting when an appropriate number of ion assignments with relatively high DP were selected (Figure 2). Since Support Vector Machine is designed in classification for binary variable, two analyses, SVM_SE (‘1’ for ‘is SE’ and ‘0’ for ‘not SE’) and SVM_ST (‘1’ for ‘is ST’ and ‘0’ for ‘not ST’), were used to target *S*. Enteritidis and *S*. Typhimurium respectively. From this we can speculate that when more serovars are targeted, additional variables for each should be created, with the potential for increased accuracy in typing.

## Discussion

MALDI-TOF mass spectra were significantly influenced by numerous experimental factors, such as the media and matrix used for bacterial growth and sample preparation, the settings of instrument (9, 24, 25), how well the technician can do his/her best to acquire just 1 µL of sample and smear it exactly the same way every time, as well as the data processing protocol for peak identification and data mining as presented in this study. It has been reported that while some ions appeared reproducible across laboratories, other unique ions were observed within laboratories when identical samples were tested using standardized protocols simultaneously at three laboratories using different instruments (26). Ion 3018 m/z specific to SE (Table 2) likely corresponds to double charges of ion 6036 m/z by Dieckmann and Malorny (5). The serovar-specific ion 7148 m/z for ST reported by Yang *et al*. (16) was not detected in our study, nor by Dieckmann and Malorny (5). We also found that the ions 10872 and 10988 m/z reported in ST (5) are shared among other serovars. Therefore, recognition of a serovar specific ion peak requires representative samples with a relatively large sample size. Decision tree-based methods would be the fastest way for typing when serovar specific biomarker(s) can be detected consistently. Unfortunately, that premise does not hold, which supports the need of more sophisticated tools in data analysis as we have carried out in this work.

Serovar identification requires different ion peaks other than the ones for species typing in MALDI-TOF MS. The ion peaks useable as biomarkers may be detected in a high or low chance. Low abundance ion peaks could be missed easily due to non-optimized instrumental settings and/or processing protocol from raw data collection to peak picking, as a result it can affect the training module dramatically in an analysis. Logarithmic transformation of intensity grants a better chance to identify the ions in low abundance. The signal/noise ratio (S/N) is theoretically proportional to sqrt(N), where N is the number of independent data sets. That’s why a reasonable replicate tests are practiced when a reference profile needs to be established or a test result to be certified. Advanced mathematical models are always required for identification of biomarkers specific to serotype(s). Inclusion of the ions with no specificity can negatively impact the discrimination. Specific and reproducible biomarkers are always preferred especially for decision tree-based models, biomarkers shared across serotypes coped with multivariate analysis and AI tools are likely the solution for *Salmonella* serotyping which requires ≥ 250 antisera to type ≥ 2600 serovars (5, 6, 20).

Successful identification of microorganisms using MALDI-TOF MS relies heavily on the database containing the spectra of known organisms (27). Identification of unknown isolates is possible only if the database contains peptide mass fingerprints of the type strains of specific genera/species/subspecies/strains (10). Confirmation via biochemical, serological and/or molecular techniques is required before including spectra for a given taxonomic unit to the database. From this perspective, databases should continue to evolve with increased number of spectra and serovars in order to discriminate among serovars more accurately. One of the practical ways then is to share data among laboratories.

A classification based on a properly established PCA model could be a useful tool for subtyping (Figure 1). The neural network and forest based deep learning tools were more sophisticated than the decision tree-based tools for *Salmonella* serotyping (Figure 2). Quality raw data, an optimized spectral processing protocol and sophisticated data mining tools incorporated into MALDI-TOF instruments or shared at research centers are likely a promising approach for rapid screening and early alert of *Salmonella* serotypes of concern and other zoonotic pathogens from the samples processed routinely in large volumes by MALDI-TOF mass spectrometry.

## Acknowledgement

This work was supported by Ontario Agri-Food Innovation Alliance, a collaboration between the Ontario Ministry of Agriculture, Food & Rural Affairs and the University of Guelph. SAS Institute Inc. kindly granted SAS licence for our academic research use.

